# In-vivo dendrite injury drives local mitochondrial contraction and dendrite branching

**DOI:** 10.1101/2025.06.05.658131

**Authors:** Patrick T Hwu, Lauren A. Kim, Marcelo A. Wood, Katherine L. Thompson-Peer

**Author notes:** Correspondence Katherine L. Thompson-Peer. Contributions Conceptualization: P.T.H. & K.T.P.; In-vivo *Drosophila* microscopy: 2-Photon Laser Injury: P.T.H.; Live imaging: P.T.H. & L.A.K.; Formal Analysis: P.T.H. & L.A.K.; Data visualization: P.T.H., K.T.P. & M.A.W.; Manuscript preparation: P.T.H., K.T.P., L.A.K. & M.A.W.; Supervision: K.T.P. & M.A.W.; Funding: K.T.P. Authors are ordered by contributions.

## Abstract

Mitochondria regulate cellular homeostasis in development and disease, and mitochondrial morphology plays a role in local injury signaling and wound repair. How mitochondria respond during dendrite injury remains an open fundamental question. Here we show that mitochondria contract rapidly and locally after laser dendrotomy. In the proximal intact dendrite, the extent of mitochondrial contraction diminishes with increasing distance from the injury site. We report that mitochondrial contraction is dependent on injury severity and that immediate contraction after injury results in a spatiotemporal increase in dendrite branching. Additionally, we find that mitochondrial contraction is inhibited by KCNJ2 (potassium inwardly rectifying channel subfamily J member 2), providing evidence that mitochondrial contraction is regulated by electrical activity. Mechanistically, we find that injury-induced mitochondrial contraction requires Drp1 (Dynamin related protein 1). In conclusion, these in-vivo findings characterize a dendrite response for mitochondria in neurons and provide insight into the regenerative outcomes of dendrites after injury.

**Graphical abstract:** In-vivo dendrite injury drives local mitochondrial contraction and dendrite branching

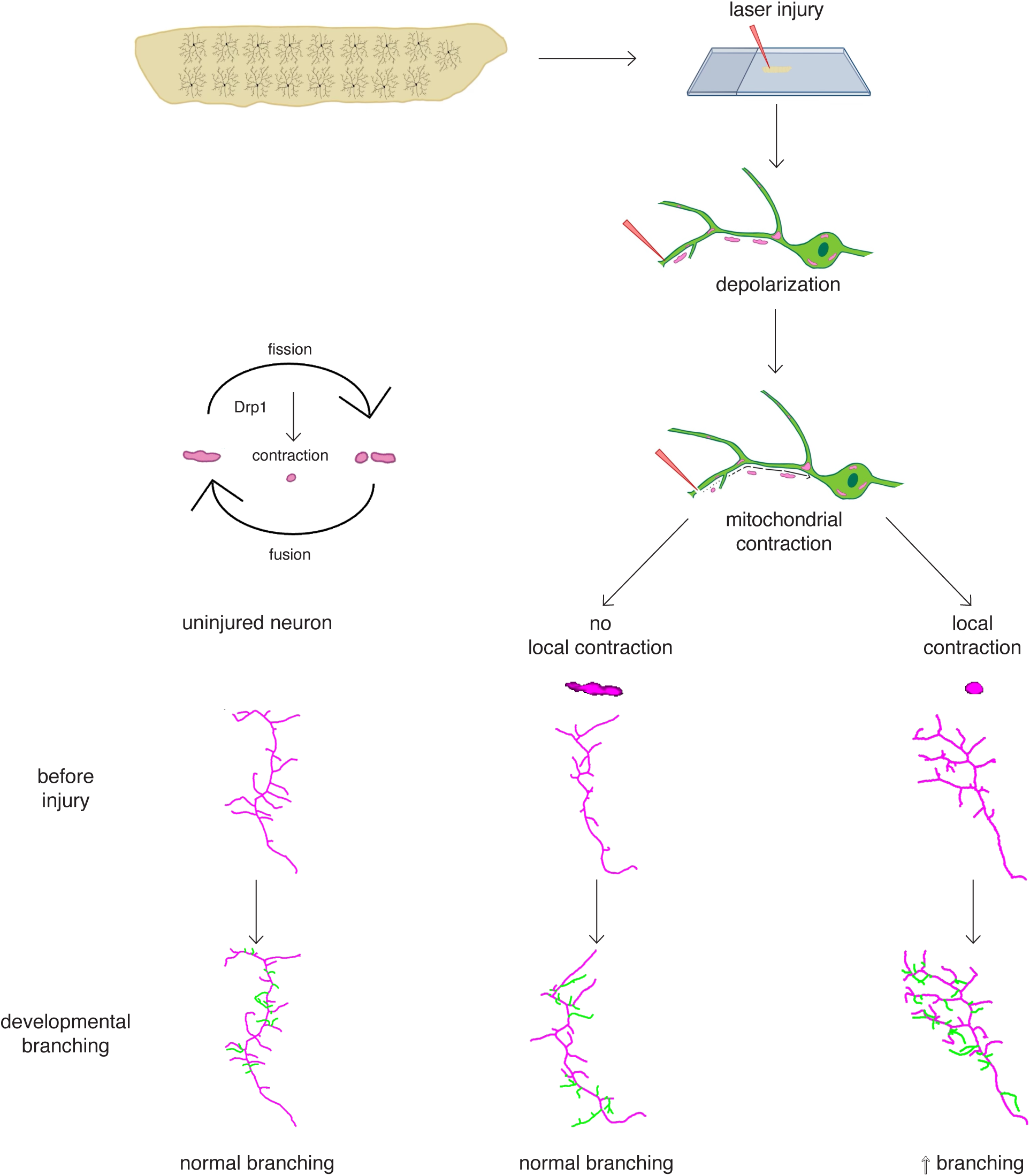

## Introduction

Cells constantly encounter mechanical and chemical injuries that trigger adaptive or pathological intracellular responses. Successful cellular repair relies on precise spatiotemporal localization of injury signals and detection both detecting abnormal physiology^1–9^. Neuronal homeostasis is disrupted in many conditions, from normal aging^10,11^ to neurodegeneration^12^, and physical injuries like spinal cord^13^ and traumatic brain damage^14^. Central to these conditions is disrupted mitochondrial homeostasis, resulting in decreased ATP production, excessive calcium buffering, and increased ROS (reactive oxygen species) production^15^. Thus, studying changes in mitochondrial integrity is critical for understanding how neurons are jeopardized during injury and disease.

Mitochondrial morphology is essential for maintaining local ATP production, protein and calcium homeostasis, and stress responses^12,16–18^. To sustain proper function, mitochondria are continuously remodeled by fusion and fission machinery, which facilitates the recycling of damaged components and removal (mitophagy) of dysfunctional organelles^19^. Fusion is regulated by the interaction of MFN1 and MFN2^20^ on the outer mitochondrial membrane, while OPA1^21^and its protease OMA1^22^ mediate inner membrane fusion. In contrast, fission is primarily driven by the translocation of DRP1 from the cytosol to the outer mitochondrial membrane, where its activity is regulated by OMM-anchored receptor proteins^23,24^. This dynamic process is critical for cellular function under basal conditions, and under stress, adopt adaptive roles, such as buffering calcium and inducing reversible morphological changes^1,2,5^. These acute adaptations influence whether a cell undergoes apoptosis or initiates repair. Despite the importance of mitochondria in injury signaling, the mechanisms regulating mitochondrial dynamics after neuronal injury and subsequently repair, are incomplete.

Neuronal injury is known to induce microtubule catastrophe, calcium influx, and changes in mitochondria morphology. Within seconds of axotomy, intracellular Ca^2+^ is upregulated, microtubules are destabilized, organelle homeostasis is disrupted, and mROS (mitochondrial reactive oxygen species) are produced^25–30^. After axon injury, mitochondria localize to the injured axon to support regeneration^31,32^. Whether dendrite injury triggers changes in mitochondrial morphology, and more criticality repair, has yet to be elucidated.

Here, we investigate mitochondrial behavior following laser dendrotomy. In vivo time-lapse imaging reveals that mitochondria in the intact dendrite rapidly contract before stabilizing, as a function of injury distance. We find that contraction scales with injury severity independent of cytoskeletal disruptions or calcium buffering. We show that disrupting neuronal activity or mitochondrial fission attenuates injury-induced mitochondrial contraction. Furthermore, we observe spatiotemporal increases in dendrite branching associated with mitochondrial contraction. These findings suggest that mitochondrial contraction may be a critical trigger for repair and not merely a byproduct of injury.

## Results

### Laser injury induces local changes in mitochondrial morphology

To assess how neuronal mitochondria respond to injury, we imaged mitochondria in class IV *Drosophila* PNS sensory neurons in live animals, before and after injury ^33,34^. We adapted LarvaSPA, an immobilization technique that preserves respiration during laser injury, because hypoxia can alter mitochondria morphology (see materials and methods)^35,36^. Axotomy (axon injury) is known to trigger a global change in the neuronal cytoskeleton (**Movie S1**) and alter mitochondrial morphology in the injured axon^29,37^. Based on this we hypothesized that axotomy may induce changes in mitochondria morphology in dendrites.

First, we tested how axotomies affected mitochondria morphology in uninjured dendrites (**Fig. 1A**). Before axotomy, we observed a mix between tubular and spherical mitochondria. After axotomy, the mitochondria in the reimaged dendrites retained their tubular morphology ∼2 mins after injury (**Fig. 1, B and C**). Next, we examined if dendrotomy (dendrite injury) alters dendritic mitochondria. Before injury, primary dendrites also had a mix of tubular and spherical mitochondria. However, upon reimaging primary and nearby attached dendrites ∼2 mins after dendrotomy, tubular mitochondria from these dendrites became spherical and lost a statistically significant amount of area (**Fig. 1B**). While the number of mitochondria fluctuated slightly, the overall quantity remained consistent after injury (**Fig. 1D**).

**Fig. 1.**
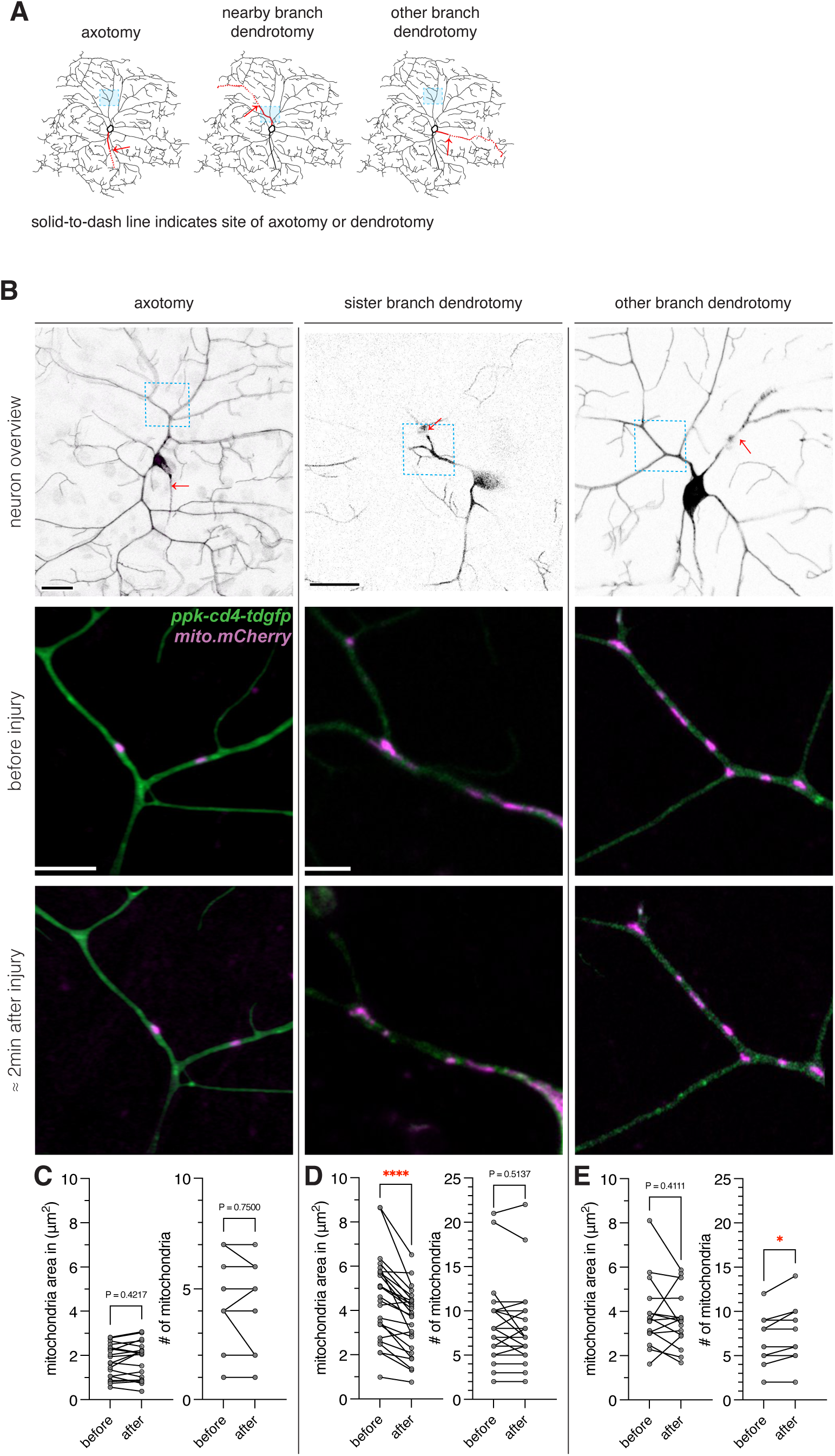
Dendrite injury induces local changes in mitochondrial morphology. (**A**) Schematic of 2-photon laser injury location (red arrow) and imaged mitochondria (blue box). (**B**) Overview of neuron before axotomy, nearby branch dendrotomy, and other branch dendrotomy (top row: left to right; red arrow). Representative image of mitochondria labeled with an outer mitochondrial membrane marker (mCherry.OMM) (magenta) in neurons (green) before axotomy, before nearby branch dendrotomy, and before other branch dendrotomy (center row: left to right). Representative image of mitochondria approximately 2 mins after axotomy, nearby branch dendrotomy, and other branch dendrotomy. Overview scale bar 40 μm, inset scale bars 5 μm. (**C**) Area and number of mitochondria in uninjured dendrites before and after axotomy. (**D**) Area and number of mitochondria in injured and surrounding dendrites before and after nearby dendrotomy. (**E**) Area and number of mitochondria area and before and after other branch dendrotomy. **N** = # of neurons, **n** = # of mitochondria analyzed. wildtype axotomies **N = 22**, **n = 75**; wildtype nearby branch dendrotomy **N = 25, n = 209**; wildtype other branch dendrotomy **N = 15, n = 100** *p<0.05, **p<0.01, ***p <0.001, ****p<0.0001, non-significant p-values are reported. Two-tailed paired t tests were used unless specified otherwise.

We sought to determine whether these mitochondrial changes were local or global. To test this, we imaged mitochondria in an uninjured primary dendrite before and after dendrotomy to a separate primary dendrite in the same neuron (other branch dendrotomy). When reimaging the uninjured dendrite, we found that mitochondria retained their tubular morphology after we dendrotomized a different primary branch (**Fig. 1E**). In summary, we find that dendrotomy triggers local changes in mitochondrial morphology.

### Dendrotomy induces distance dependent mitochondrial contraction

Our observations suggested that dendrotomy induces mitochondrial contraction, a phenomenon that is observed in the axon after axotomy ^26^. We hypothesized that mitochondria respond to laser dendrotomy locally, because we primarily observed changes in mitochondrial area but not quantity. To interrogate how dendrotomy induced changes in mitochondria morphology, we conducted live-imaging on mitochondria during proximal and distal primary dendrotomies (**Fig. 2A**).

**Fig. 2.**
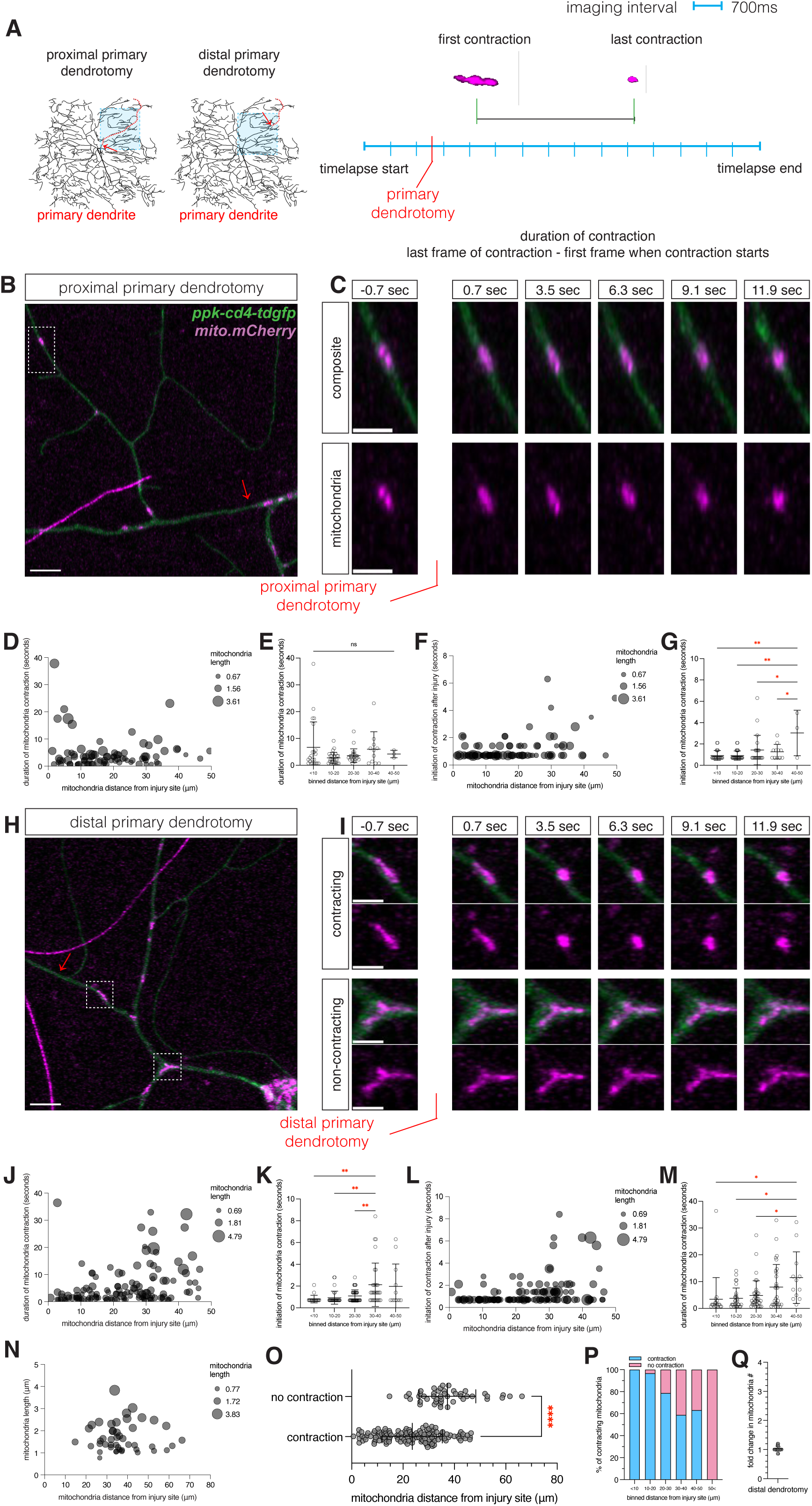
Injury induced mitochondrial contraction is dependent on distance from injury site. (**A**) Schematic of injury location for proximal primary or distal primary dendrotomies. Images of neurons were continuously acquired for 200 frames and dendrites were injured at approximately frame 10. (**B**) Representative proximal primary dendrotomy live imaging and dendrotomy location (red arrow). Scale bar 5 μm. (**C**) Magnified view of mitochondrial contraction immediately ∼0.700 msec before and after dendrotomy. Scale bar 2 μm. (**D**) Characterization of mitochondrial contraction in the severed dendrite after primary proximal dendrotomy. (**E**) Duration of mitochondrial contraction after proximal primary dendrotomy binned by distance from dendrotomy site, N.S. (**F**) Duration before mitochondria initiated contraction after proximal primary dendrotomy relative to mitochondrial length. (**G**) Duration before mitochondria initiated contraction after proximal primary dendrotomy (<10 μm vs. 40-50 μm, p=0.0026**; 10-20 μm vs. 40-50 μm, p=0.0022**, 20-30 μm vs. 40-50 μm, p=0.0479*; 30-40 μm vs. 40-50 μm, p=0.0345*, One-way ANOVAs, Bonferroni corrected) binned by distance from the dendrotomy site. (**H**) Representative distal primary dendrotomy live imaging and dendrotomy location (red arrow) areas of mitochondrial contraction (‘white box) and no mitochondrial contraction (“white box). Scale bar 5 μm. (**I**) Magnified view of contracting (top panel) and non-contracting (bottom panel) mitochondria ∼0.700 msec before and after injury. Scale bar 2 μm. (**J**) Characterization of mitochondrial contraction in proximal intact dendrites in neurons after distal primary dendrotomy. (**K**) Duration of mitochondrial contraction after distal primary dendrotomy (<10 μm vs. 40-50 μm, p=0.0184*; 10-20 μm vs. 40-50 μm, p=0.0132*, 20-30 vs. 40-50 μm, p=0.0405*, One-way ANOVAs, Bonferroni corrected) binned by distance from dendrotomy site. (**L**) Duration before mitochondria initiated contraction after distal primary dendrotomy relative to mitochondrial length. (**M**) Duration before mitochondria initiated contraction (<10 μm vs. 30-40 μm, p=0.0040**; 10-20 μm vs. 30-40 μm, p=0.0034**, 20-30 μm vs. 30-40 μm, p=0.0093**, One-way ANOVAs, Bonferroni corrected) binned by distance from dendrotomy site. (**N**) Distance of individual mitochondria from the dendrotomy site that failed to contract in the intact dendrite after distal primary dendrotomy. (**O**) Mean distance of contracting vs non-contracting mitochondria after distal primary dendrotomy. (**P**) Percent of contracting (blue) vs non contracting (pink) mitochondria binned by distance from dendrotomy site: 19/19 contraction <10 μm; 29/30 contraction 10-20 μm; 37/47 contraction 20-30 μm; 30/51 contraction 30-40 μm; 12/19 contraction 40-50 μm; 0/7 contraction >50 μm. (**Q**) Fold change in number of mitochondria after distal primary dendrotomy. **N** = # of neurons, **n** = # of mitochondria analyzed. wildtype proximal dendrotomies **N = 20, n = 81**; wildtype distal dendrotomies **N = 30, n = 173** Data presented as mean ± SD. *p<0.05, **p<0.01, ***p <0.001, ****p<0.0001. Two-tailed Mann-Whitney test, unless specified.

First, we aimed to understand how injury alters mitochondrial morphology in the distal severed dendrite during proximal primary dendrotomies. Before proximal primary dendrotomy, primary dendrites are decorated with tubular mitochondria. However, immediately after dendrotomy tubular mitochondria contract rapidly and completely in the distal severed segment (**Fig. 2, B and C; Movie S2**). Furthermore, 84 out of 84 mitochondria contracted independent of distance from dendrotomy site, mitochondrial length, and contraction duration (**Fig. 2, D and E**). However, mitochondria initiated contraction later as the distance between the dendrotomy site and individual mitochondria increased (**Fig. 2, F and G**).

We next examined how mitochondria respond in the proximal intact dendrite–which remains attached to the cell body–after distal primary dendrotomy (**Fig. 2H; Movie S3**). Immediately after dendrotomy, mitochondria in the proximal intact dendrite contracted rapidly (**Fig. 2, I’ and I’’**). Furthermore, we found that mitochondria contracted faster and were more likely to contract if they were closer to the dendrotomy site (**Fig. 2, J and K**). Furthermore, we observed that mitochondria closest to the injury site initiated contracted earlier than mitochondria furthest from the injury site (**Fig. 2, L and M**). While we observed mitochondrial contraction in 100% of the neurons we injured (30 out of 30), not every mitochondria in each neuron contracted; mitochondria were less likely to contract if they were further than 30 *μ*m away from the dendrotomy site (**Fig. 2, N and O**). We observed that mitochondria within <20 *μ*m of the dendrotomy site contracted in majority of cases (48 out of 49), whereas mitochondria further than 50 *μ*m from the dendrotomy site never contracted (**Fig. 2P**). Moreover, we validated that mitochondria were primarily contracting rather than undergoing fission by counting the number of mitochondria before and after dendrotomy (**Fig. 2Q**).

To compare how axotomy affects mitochondrial morphology in the injured axons versus what we observed in dendrites, we performed live-imaging of mitochondria in axons during axotomy (**Fig. S1A**). Similar to the mitochondria morphology we observed in dendrites, axonal mitochondria contain a mix of tubular and spherical morphologies before axotomy (**Fig. S1B**). Immediately after axotomy, 78 out of 78 of mitochondria in the intact axon contracted rapidly and completely (**Fig. S1, C and D; Movie S4**). We noted a delay in contraction initiation and duration as the distance between axotomy site and mitochondria increased (**Fig. S1, E and F)**. In summary, we find that duration and timing of injury induced mitochondrial contraction is dependent on distance from the injury site in the proximal intact dendrite.

### Mitochondria contract independently of microtubule catastrophe or mitochondrial calcium buffering

Mitochondrial morphology is partly regulated through interactions with microtubules ^38^. To determine whether injury-induced microtubule catastrophe drives mitochondrial contraction we conducted distal primary dendrotomies while live-imaging both microtubules (using the *plus-end binding protein, EB1-GFP*) and mitochondria. Indeed, we observed changes in microtubules–evident by the sudden assembly of EB1-GFP puncta in dendrites–immediately after dendrotomy. In 72.4% (21 out of 29) of the neurons, mitochondrial contraction and EB1 puncta formation occurred simultaneously (**Fig. S2A; Movie S5**). However, there were instances when microtubule disruption and mitochondrial contraction were not correlated: EB1-GFP puncta formed prior to mitochondrial contraction (∼10.34%), EB1-GFP puncta were present without mitochondrial contraction (∼3.45%) (**Movie S6**), mitochondria contracted prior to EB1-GFP puncta forming (∼6.90%), and mitochondrial contraction occurred in the absence of EB1-GFP puncta formation (∼6.90%) (**Fig. S2B; Movie S7**). In summary, we find that mitochondria contract independently of microtubule catastrophe.

Both axotomies and dendrotomies are known to trigger a surge of cytosolic calcium influx ^39,40^. In response to elevated intracellular calcium, mitochondria rapidly buffer calcium to restore calcium back to baseline levels ^41^. Furthermore, mitochondrial calcium buffering is known to affect mitochondrial morphology ^42^. We hypothesized that mitochondrial calcium buffering promotes mitochondrial contraction. To test this, we performed distal primary dendrotomies and tertiary dendrotomies (tertiary dendrite injury) and imaged Ca^2+^_m_ (mitochondrial calcium) levels using a mitochondrial GCaMP marker (**Fig. S3A**). Before distal primary dendrotomies, mitochondria are primarily undetectable with the mitoGCaMP marker. However, following distal primary dendrotomy, we observed a retroactive wave of Ca^2+^_m_ uptake–originating from the dendrotomy site that propagated to uninjured dendrites, through the cell body, and finally down the axon (**Fig. S3, A and A’; Movie S8)**. We observed an increase in Ca^2+^_m_ levels and contraction in the mitochondria near the dendrotomy site. Despite the increase in Ca^2+^_m_ in uninjured dendrites and the cell body, we failed to observe mitochondrial contraction in the uninjured dendrites or in the cell body (**Fig. S3, A’’ and A’’’, Fig. S3B**). In summary we find that sudden mitochondrial calcium influx is not the cause of injury-induced mitochondrial contraction.

### Injury-induced mitochondrial contraction scales with dendrotomy severity

Injury-induced changes in mitochondrial morphology are known to activate the injury signaling cascade ^1,2,5^. In order to evaluate the consequences of mitochondrial contraction, we designed a paradigm that allowed us to injure neurons but induce contraction in only a subset of injured neurons.

To investigate the consequence of mitochondrial contraction, we imaged the neuronal architecture (before dendrotomy), conducted live-imaging of tertiary dendrotomies (tertiary dendrite injury), then reimaged the neuronal architecture 24 hrs after dendrotomy (**Fig. 3A**). We confirmed that tertiary dendrotomies were successful by the presence of a physically severed branch (**Fig. 3B, Fig. S4A; Movie S9 and Movie S10**). After tertiary dendrotomy, nearby mitochondria contracted in some neurons (**Fig. 3C**) which we confirmed by counting the number of mitochondria before and after dendrotomy (**Fig. 3D**). Furthermore, we noticed that mitochondria failed to contract if the distance between mitochondria and the dendrotomy site exceeded 30 *μ*m (Fig. 3E). We observed that tertiary dendrotomies were less likely to induce mitochondrial contraction as only 38 out of 103 mitochondria contracted (**Fig. 3, E-G, Fig. S4B**). Consistent with our observations after primary dendrotomy, mitochondria closer to the dendrotomy site were more likely to contract. However, even at similar distances away from the dendrotomy site, mitochondrial contraction after tertiary dendrotomy expended more time contracting (**Fig. S4C**).

**Fig. 3.**
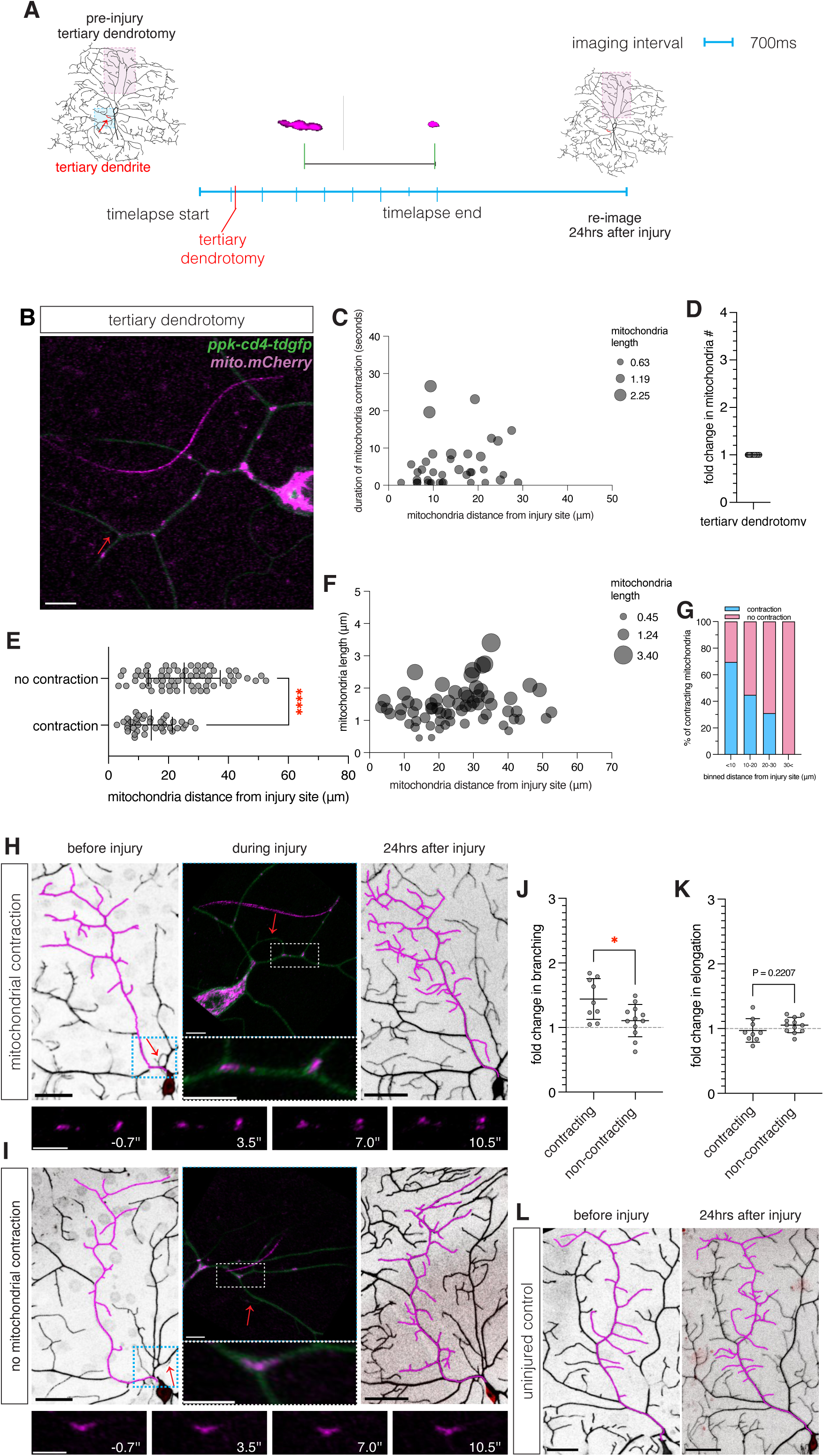
Injury-induced mitochondrial contraction is associated with a spatiotemporal increase in dendrite branching. **(A**) Schematic of tertiary dendrotomy. An overview of the neuron architecture was taken before dendrotomy. Images of neurons were continuously acquired for 200 frames and dendrites were injured at approximately frame 10. Neurons were imaged 24 hrs. (**B**) Representative live imaging of tertiary dendrotomy and dendrotomy location (red arrow) and locations of contracting (‘white box) vs non-contracting (“white box) mitochondria. Scale bar 5 μm. (**C**) Magnified view of contracting mitochondria (top panel) vs non contracting mitochondria (bottom panel) ∼0.700 msec before injury and after injury. (**D**) Fold change in number of mitochondria after tertiary dendrotomy. (**E**) Percent of contracting (blue) vs non-contracting (pink) mitochondria binned by distance from dendrotomy site: 16/23 contraction <10 μm; 13/29 contraction 10-20 μm; 9/29 contraction 20-30 μm; 0/22 contraction 30 μm<. (F) Characterization of mitochondrial in proximal intact dendrites after tertiary dendrotomy. (**G**) Distance of individual mitochondria from the dendrotomy site that failed to contract in relation to mitochondrial length (**H**) Mean distance of contracting mitochondria vs non-contracting mitochondria after tertiary dendrotomy. Two-tailed Welch’s t test. (**I**) Representative overview of neuronal architecture with contracting mitochondria before (left), during (middle), and 24 hrs after (right) tertiary dendrotomy. Scale bar 25 μm, 5 μm, 50 μm. (**J**) Representative overview of neuronal architecture without contracting mitochondria before (left), during (middle), and 24 hrs after (right) tertiary dendrotomy. Scale bar 25 μm, 5 μm, 50 μm. (**K**) Change in dendrite branching rate 24 hrs after tertiary dendrotomy in neurons with contracting (N=9) vs non-contracting mitochondria (N=12) (**L**) Change in dendrite elongation rate 24 hrs after tertiary dendrotomy in neurons with contracting (N=9) vs non-contracting mitochondria (N=12).(**M**) Representative image of uninjured control neuron on the day of dendrotomy and 24 hrs after. **N** = # of neurons, **n** = # of mitochondria analyzed. wildtype tertiary dendrotomy **N = 36, n = 103**; wildtype dendrotomy induced branching **N = 21** Data presented as mean ± SD. *p<0.05, **p<0.01, ***p <0.001, ****p<0.0001, non-significant p-values are reported. Unpaired two-tailed t tests, unless specified.

To test if mitochondrial contraction is associated with initiating a repair response, we took advantage of tertiary dendrotomies, as these injuries only sometimes provoked contraction (**Fig. 3E**). We imaged the entire architecture of the neuron before dendrotomy, conducted time-lapse imaging of tertiary dendrotomies, and reimaged the neuronal architecture 24 hrs after dendrotomy (**Fig. 3, H and I; Movie S11 and Movie S12**). One technical challenge with tertiary dendrotomies is that tertiary dendrites are short (<10 *μ*m) and adjacent to the shaft of the primary dendrite. To ensure that we were only assessing the consequence of mitochondrial contraction on dendrite growth, we only quantified neurons if the attached primary branch was present before and 24 hrs after dendrotomy (**Movie S13**). To account for intra-neuronal and inter-animal differences in dendrite growth, we normalized dendrite branching rates of the injured neuron to an uninjured neuron within the same animal (**Fig. 3l**). We observed that 24 hrs after tertiary dendrotomy, neurons *with* contracting mitochondria display a statistically significant increase in dendrite branching compared to neurons *without* contracting mitochondria (**Fig. 3J**); while no significant differences were detected in dendrite elongation (**Fig. 3K**). In summary, we observe that tertiary dendrotomy induces a lesser extent of mitochondrial contraction and mitochondrial contraction is associated with a spatiotemporal increase in dendrite branching.

### Electrical hyperpolarization via *KCNJ2* impairs mitochondrial contraction after primary dendrotomy

Electrical activity is a well-established regulator of dendrite patterning during development ^43^. Previous work has shown that hyperpolarizing the resting membrane potential by overexpressing *KCNJ2 (potassium inwardly rectifying channel subfamily J member 2)* inhibits dendrite regeneration ^44^, partly by blocking calcium-mediated injury detection ^40^.

We postulated that injury-induced mitochondrial contraction requires normal electrical activity. To test this, we live-imaged mitochondria during primary dendrotomies, since primary dendrotomies elicit a stronger contraction response than tertiary dendrotomies (**Fig. 4A**). Prior to dendrotomy, mitochondria in *KCNJ2* neurons are not abnormally tubular or spherical (**Fig. 4, B and C; Movie S14**). We observed that overexpressing *KCNJ2* significantly inhibited mitochondrial contraction after a primary dendrotomy (**Fig. 4D**). However, we also observed instances where *KCNJ2* expression partially suppressed injury-induced mitochondrial contraction. To better understand how *KCNJ2* inhibits mitochondrial contraction, we plotted the distance between individual mitochondria and the dendrotomy site. Our results reveal that *KCNJ2* increases the duration of mitochondrial contraction even when mitochondria are immediately adjacent to the dendrotomy site (**Fig. 4E; Movie S15**).

**Fig. 4.**
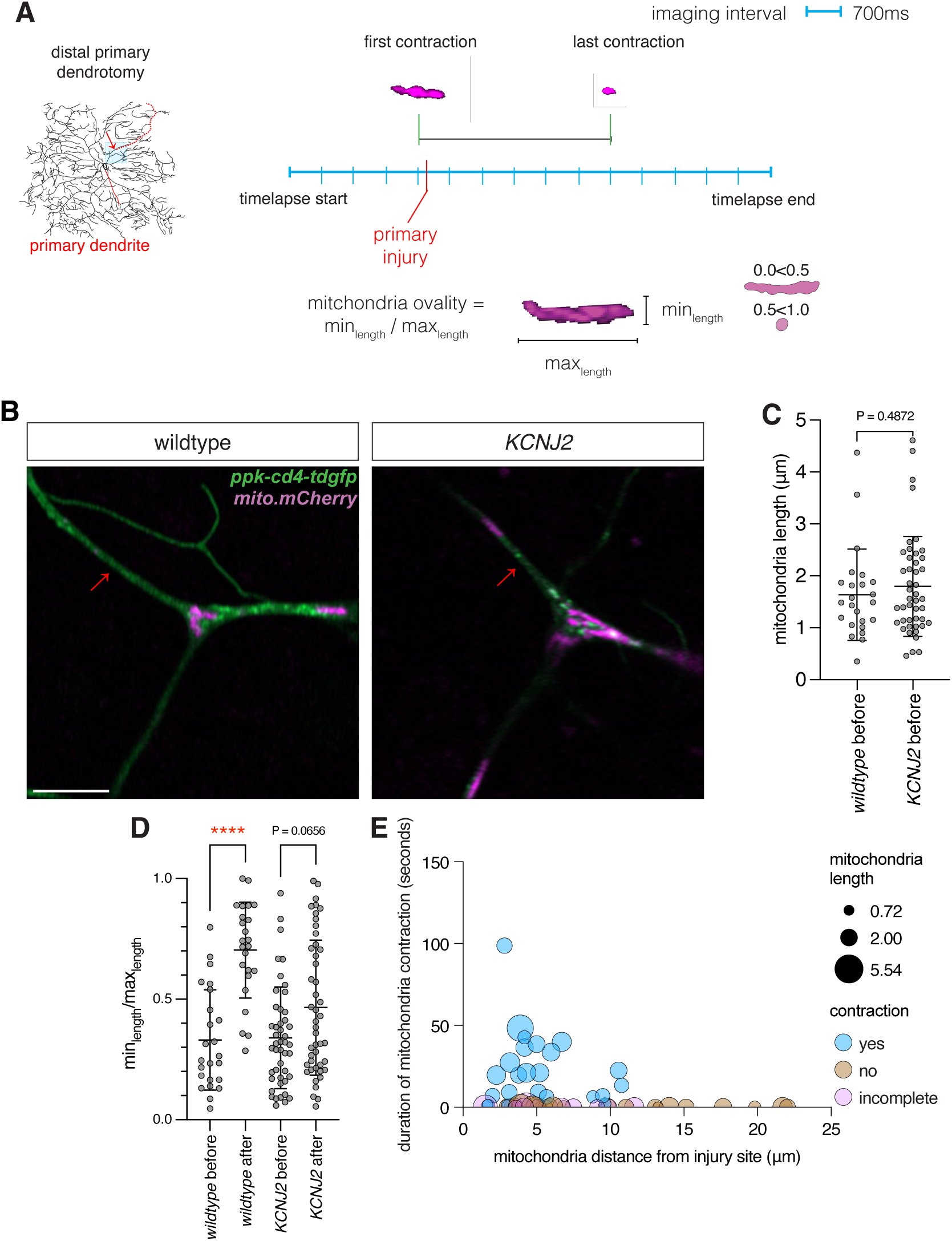
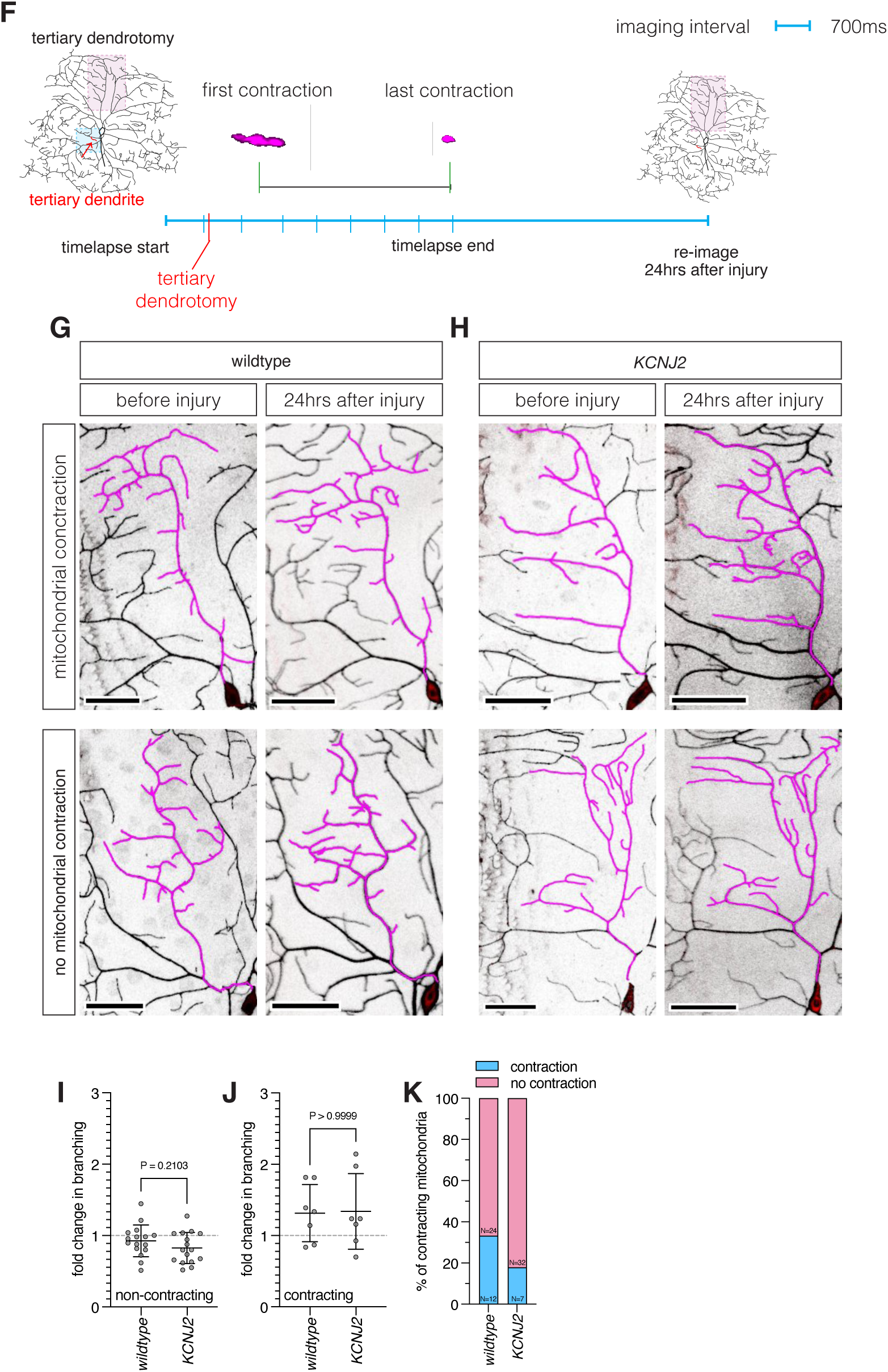
Clamping neuronal membrane potential inhibits mitochondrial contraction after primary dendrotomy. (**A**) Schematic of primary dendrotomy. Mitochondrial ovality was calculated by taking the min_length_/max_length_ of individual mitochondria before and after dendrotomy. Live images were acquired for 200 frames and neurons were injured between frame 1 and frame 20. (**B**) Representative images of neurons and dendrotomy location (red arrow). Scale bar 5 μm. (**C**) Maximum mitochondrial length before dendrotomy in wildtype vs KCNJ2 neurons. (**D**) Mitochondrial ovality before and after primary dendrotomy in wildtype neurons and KCNJ2 neurons. One-way ANOVA, Bonferroni corrected. (**E**) Contracting mitochondria in KCNJ2 after primary dendrotomy. (**F**) Schematic of tertiary dendrotomy and imaging location after dendrotomy. (**G, H**) Bottom row: wildtype versus KCNJ2 dendrite architectures for neurons that had no change in branching in the absence of mitochondrial contraction. Top row: wildtype versus KCNJ2 dendrite architectures for neurons that had dendrotomy induced branching with mitochondrial contraction. Scale bar before dendrotomy 25 μm; scale bar 24 hrs after 40 μm). (**I**) 24hr change in branching rates for neurons without contracting mitochondria in wildtype (N=16) vs KCNJ2 (N=16) after tertiary dendrotomy. Unpaired, two-tailed, t test. (**J**) 24hr change in branching rates for neurons with contracting mitochondria in wildtype (N=7) vs KCNJ2 (N=7) after tertiary dendrotomy. (**K**) Percentage of dendrotomy induced mitochondrial contraction in wildtype (N=12 contracting, N=24 non contracting) vs KCNJ2 (N=7 contracting, N=32 non contracting) after tertiary dendrotomy. **N** = # of neurons, **n** = # of mitochondria analyzed. wildtype primary dendrotomy **N = 13, n = 48**; KCNJ2 primary dendrotomy **N = 21, n = 92** wildtype tertiary dendrotomy **N =** 36; KCNJ2 tertiary dendrotomy **N = 39** Data presented as mean ± SD. *p<0.05, **p<0.01, ***p <0.001, ****p<0.0001, non-significant p-values are reported. Mann-Whitney tests, unless specified.

We extended our findings that expressing *KCNJ2* attenuates injury-induced mitochondrial contraction by live imaging *KCNJ2* neurons during tertiary dendrotomies (**Fig. 4, F-H**). During live imaging, we observed that injury-induced mitochondrial contraction in wildtype and *KCNJ2* expressing animals occurred in a stochastic nature. To better understand if *KCNJ2* expressing animals displayed differences in the dendrite architecture, we tracked the dendrite arbor before and after dendrotomy. In wildtype and *KCNJ2*, neurons without contracting mitochondria, we failed to observe an increase in dendrite branching compared to their uninjured counterparts (**Fig. 4I**). Furthermore, we observed increased dendrite branching in the presence of contracting mitochondria in wildtype and *KCNJ2* neurons (**Fig. 4J**). Lastly, we observed less instances of injury-induced mitochondrial contraction after tertiary dendrotomy in *KCNJ2* (7/39, ∼17.95%) compared to wildtype neurons (12/36, ∼33.33%) (**Fig. 4K; Movie S16 and Movie S17**). In summary, we find that injury-induced mitochondrial contraction requires normal neuronal activity.

### Complete mitochondrial contraction requires *Drp1* activity

Previous studies have shown that injury-induced mitochondrial fragmentation is regulated through *Drp1* (*Dynamin related protein 1*)^7,30,45–47^. We hypothesized that injury-induced mitochondrial contraction requires *Drp1*. To test this, we conducted distal primary dendrotomies in GTPase dead *Drp1* expressing neurons, caused by a single residue (*Drp1.K38A*) substitution in the G1 motif in the GTPase domain.^48^ Before injury, *Drp1.K38A* neurons primarily have a single continuous hyperfused mitochondria (**Fig. 5A**). Immediately after distal primary dendrotomy, wildtype neurons contract rapidly and completely. However, upon distal primary dendrotomy in *Drp1.K38A* neurons, mitochondria failed to completely contract (**Fig. 5B**; **Movie S18**). We counted the number of tubular and spherical mitochondria before and after injury and observed that *Drp1.K38A* inhibited injury-induced contraction (**Fig. 5, C and D**). In summary, these findings show that complete mitochondrial contraction requires *Drp1*.

**Fig. 5.**
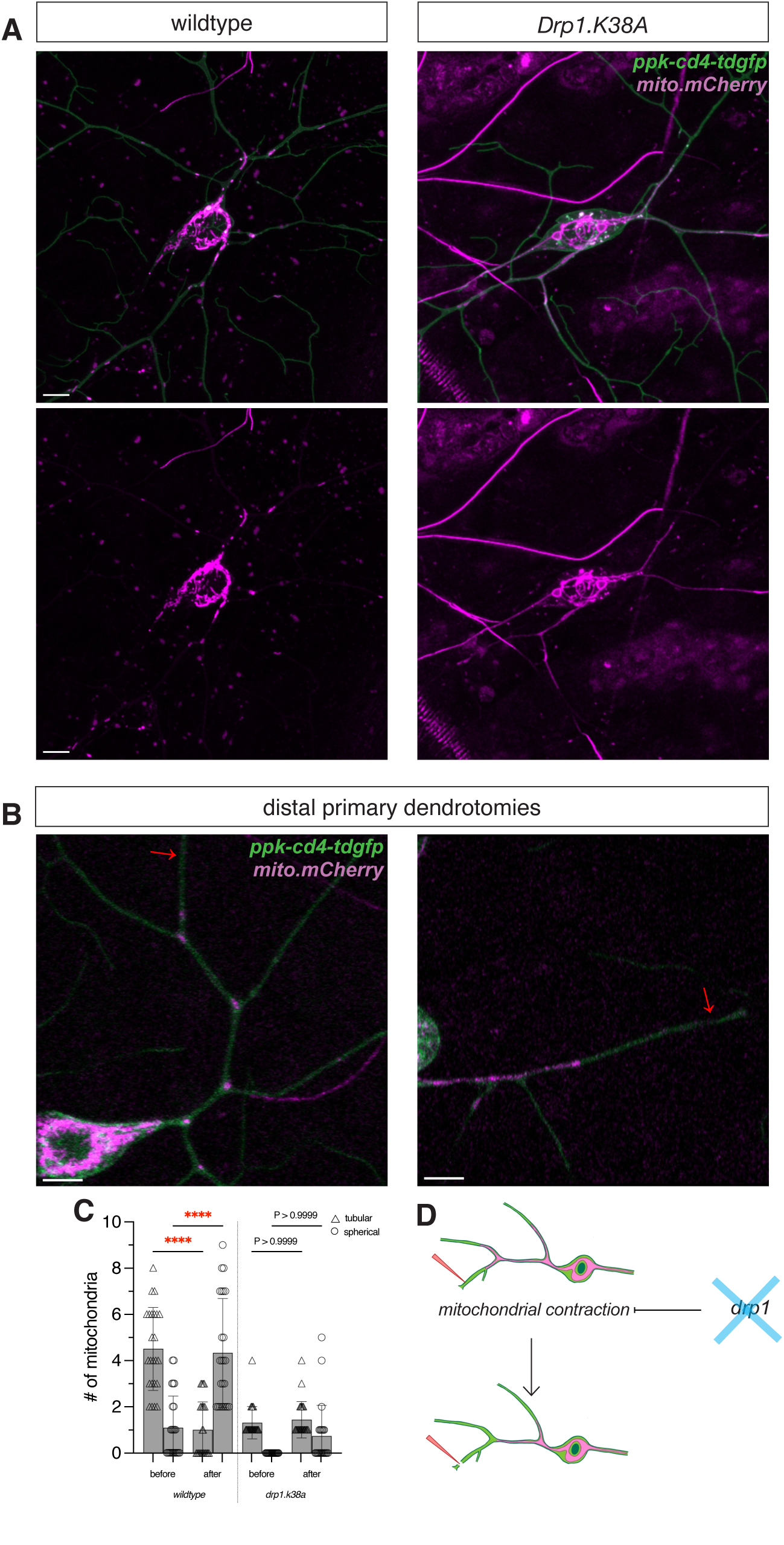
Mitochondrial contraction requires Drp1 activity. (**A**) Overview hyperfused mitochondrial network in wildtype vs Drp1.K38A expressing animals. Scale bar 5 μm. (**B**) Snapshot of ROI dendrotomy location (red arrow). (**C**) Quantification of total number of tubular and spherical mitochondria before and after dendrotomy (wildtype before tubular vs wildtype after tubular, p<0.0001****, wildtype before punctate vs wildtype after punctate, p<0.0001****). Scale bar 5 μm. (**D**) Graphical representation of inhibited mitochondrial contraction without Drp1 activity. **N** = # of neurons, **n** = # of mitochondria analyzed. Data presented as mean ± SD. *p<0.05, **p<0.01, ***p <0.001, ****p<0.0001, non-significant p-values are reported. wildtype dendrotomy **N = 24**; Drp1.K38A dendrotomy **N = 23** One-way ANOVAs, Bonferroni corrected unless specified.

## Discussion

While the role of mitochondrial injury signaling and repair is well established, the role in injury signaling and repair has yet to be elucidated in neurons, and specifically dendrites. Using our model organism’s capacity for longitudinal imaging, we monitored how mitochondria respond in real-time during dendrotomy (dendrite injury), reimaging the same dendrotomized neurons to assess repair. In this study, we highlight the spatiotemporal dynamics of mitochondrial contraction following dendrotomy. Our findings reveal that in the proximal intact dendrite, mitochondrial contraction depends on the distance from the dendrotomy site, influencing both its timing and duration. In contrast, within the severed dendrite, only the initiation of contraction is distance-dependent. Additionally, tertiary dendrotomies elicit a weaker contraction response than primary dendrotomies; at equivalent distances from the injury site, a milder injury prolongs the time to full contraction.

Given the stochastic nature of mitochondrial contraction, we examined its functional consequences. To dissect the relationship between mitochondrial contraction and repair, we inhibited dendrite repair by clamping the resting membrane potential with *KCNJ2* and assessed its impact on mitochondrial contraction. After primary and tertiary dendrotomy, mitochondrial contraction was markedly suppressed. Interestingly, *KCNJ2* expressing neurons still exhibited increased dendrite branching in the presence of injury-induced mitochondrial contraction, providing insight into the relationship between injury induced mitochondrial contraction and repair. Furthermore, we observe that injury-induced mitochondrial contraction is regulated by *Drp1* and expressing a dominant negative allele of *Drp1* inhibits complete mitochondrial contraction. Our findings are supported by previous studies demonstrating that injury induces immediate changes in mitochondrial morphology in the epidermis, MEFs, HeLa, HEK293t, c2c12^1,2,5,49,50^. Together our findings suggest that mitochondrial contraction may act as a critical trigger for dendrites to initiate repair and not a byproduct of injury.

Cellular repair is strongly influenced by the magnitude of injury: severe damage often leads to cell death, whereas milder injuries can elicit adaptive responses. In this study, we observed that mitochondria undergo immediate contraction following dendrotomy, and the extent of this contraction scales with injury severity (primary vs. tertiary dendrotomy). Our findings are consistent with previous work in live mouse axons, where electrical nerve stimulation and, more severely, axotomy, induced distance-dependent mitochondrial contraction^26^. We built upon this established framework by tracking individual neuronal responses and the consequences of dendrite repair. Future studies should build upon our findings to determine whether mitochondrial contraction serves as a direct trigger for regenerative programs or reflects a broader cellular adaptation to injury.

While the phenotypic changes in mitochondrial morphology following injury is well established, the mechanisms governing this response are only partially understood. In neurons, the fact that mitochondria contract in both the axon and dendrites highlights the conserved nature of injury-induced mitochondrial remodeling^30^. Our findings that *KCNJ2* or *Drp1.K38A* both suppress injury-induced mitochondrial contraction provides insight into previous literature; previous reports have shown that expressing *KCNJ2*^51^ or *Drp1.K38A*^52^ prevents cell death after injury. Whether KCNJ2 is involved in regulating injury-induced changes in mitochondrial morphology has yet to be elucidated until now. Future studies should determine whether *KCNJ2* and the *Drp1* converge on a single molecular pathway or diverge.

Downstream of morphological changes in mitochondria are mROS (mitochondrial reactive oxygen species). Previous studies have shown that mROS become upregulated in response to cell injury across various tissues^2,3,53–56^. Similarly, mROS is upregulated as a consequence of neuronal activity^57,58^ caused by spreading depolarization^26,59^. Lastly, axotomy induced mitochondrial contraction has been shown to stimulate mROS production^26^. Whether mROS functions downstream of mitochondrial contraction after dendrotomy and potentially a primary driver of dendrite repair remains an open question. Whether injury-induced mitochondrial contraction is directly linked to the repair process requires further investigation. Collectively our findings highlight an intricate relationship between mitochondrial dynamics and dendrite repair, emphasizing mitochondrial contraction as a potential mediator of injury-induced repair and the need for further investigation into the molecular mechanisms that govern this process.

## Material and Methods

### Animals

#### Fly Husbandry and genetics

Adult flies were reared at ∼22.5°C and fed a standard yeast diet.

#### Drosophila larvae collection and handling

Male and female *Drosophila* larvae were collected from synchronized egg lays on 4.0% grape agar plates (750ml H_2_O; 18.75g BD Difco^TM^ Agar, Mfr. No. 214010; 250ml Welch’s 100% grape juice; 25g sucrose; 1.5g methyl p-hydroxybenzoate, ReagentPlus^®^, Mfr. No. H5501-100G) supplemented with yeast paste (made from active dry yeast + dH_2_O). The plates were maintained at room temperature (22.5°C), and larvae were aged for 72 hours after egg laying (AEL).

For regeneration experiments, individual larvae were transferred to petri dishes containing 4.0% grape agar and yeast paste after injury. Before imaging, larvae were washed in distilled water (dH₂O).

#### Larval immobilization for imaging

Early third instar larvae were transferred to glass wells containing dH_2_O to wash off yeast. Class IV (ddaC) *Drosophila* PNS sensory neurons^33^ were imaged by immobilizing larvae dorsally on a ∼4.0% gelatinous agarose pad (prepared using dH₂O and LE Quick Dissolve Agarose, GeneMate, Cat. No. E3119-500), with a small volume of glycerol (Glycerol, ReagentPlus^®^, Mfr. No. G7757-500ML) which was sandwiched between a microscope slide (Fisherbrand, Cat. No. 125441, 75 × 25 × 1 mm) and a No. 1.5 coverslip (Epredia, Cat. No. 22-050-218, 22 × 22 mm) as previously described^60^.

Late third-instar larvae were immobilized using the same method, with the addition of vacuum grease (DuPont Molykote®, Electron Microscopy Sciences, Cat. No. 60705) and Scotch® tape. This immobilization method was used to generate data for **Figures 2, S2, 3, S3, 4F–4K, S4, 5**.

Larvae were immobilized using the LarvaSPA method, as previously described^35^, with ice as a temporary anesthetic instead of isoflurane. This immobilization method was used to generate data for **Figures 1C–1E, 4C–4E**.

### Microscopy

#### Confocal imaging

Confocal imaging on the Zeiss LSM980 was used to acquire static images of the dendrite architecture before injury and immediately after injury. A Zeiss LSM700 confocal microscope equipped with a 488 nm and 561 nm laser line was used to acquire static images 24hrs after injury using a ×20/0.8 N.A. air objective.

#### Airyscan imaging

A Zeiss LSM900 equipped with a 488 nm and 561 nm laser line was used to acquire static images of mitochondrial morphology before and after injury. The imaging settings are as follows: ×63/1.4 N.A. oil-immersion objective, with an imaging speed of ∼4.63 Hz and a 28.80 × 28.80 *μ*m frame size (∼216.19 ms frame time, ∼3.05 *μ*s pixel dwell time).

#### Live imaging

Live imaging was performed by collecting z-stacks using Airyscan imaging or manual z-plane focusing with regular laser scanning confocal microscopy over 200 cycles using a ×63/1.4 N.A. oil-immersion objective. In the cases of z-drift during z-stack acquisition, imaging was paused, refocused, and resumed.

#### 2-Photon laser injury

A Zeiss LSM780 and LSM980 inverted microscopes, equipped with 488 nm and 561 nm laser lines and a tunable Ti:Sapphire 2-photon laser (Mai Tai, Spectra-Physics), were used for laser injuries. Early third instar larvae were immobilized using the methods described above. For proximal primary dendrotomies, distal primary dendrotomies, and tertiary dendrotomies, a Zeiss LSM980 with a Z-piezo stage was used to image neurons and mitochondria and to perform injuries in third-instar Drosophila larvae, as previously described^61^.

On the Zeiss LSM980, injuries were performed using a ×63/1.4 N.A. oil-immersion objective, with an imaging speed of ∼11.43 Hz and a 44.9 × 44.9 *μ*m frame size (∼75.97 ms frame time, ∼0.46 *μ*s pixel dwell time, 200 cycles). A 0.4 × 0.4 *μ*m ROI, focused on a dendrite, was injured using a laser at 820 nm with a measured power of ∼84.1 mW for 1 iterations. For 400 ms imaging and injury parameters, injuries were performed using a ×63x/1.4 N.A. oil-immersion objective, with an imaging speed of ∼8 Hz and a 22.4 × 22.4 *μ*m frame size (∼121.65 ms frame time, ∼1.49 *μ*s pixel dwell time, 200 cycles). A manually controlled 0.4 × 0.4 *μ*m ROI was positioned over an axon and injured using a laser at 820 nm with a measured power of 101.5 mW for 6 iterations.

For injury-induced branching experiments two neurons were used per animal, one injured and one uninjured. For the injured neuron, a z-stack was acquired of the entire neuron architecture before conducting live-imaging of dendrotomy. We ablated neurons between and adjacent to injured and uninjured neurons (i.e. ablate - A1; dendrotomy - A2; ablate - A3; uninjured control - A4; ablate - A5).

On the Zeiss LSM780, a 0.5 × 0.5 *μ*m ROI was created over an axon or dendrite and injured for ∼2 sec at 915 nm with a measured power of ∼79.6 mW.

#### Microtubule imaging

Homozygous EB1-GFP animals were immobilized and injured using the techniques described above. A single Z-plane was acquired with an imaging speed of ∼1.43 Hz. The imaging frame was manually adjusted for focus.

#### MitoGCaMP imaging

Heterozygous MitoGCaMP animals were immobilized and injured using the parameters techniques described above.

### Image analysis

All quantifications were conducted using the open source software: ImageJ/Fiji (https://fiji.sc).

#### Mitochondrial contraction

Mitochondrial distance from the injury site was measured in ImageJ/Fiji using the segmented line tool to trace the dendritic length from the injury site to each mitochondrion within the proximal intact dendrite. Contraction onset was defined by the first morphological shift from a tubular to a circular structure following 2-photon laser dendrotomy or axotomy. The end of contraction was marked by the first frame in which mitochondria appeared fully circular, and contraction duration was calculated as the time between initiation and completion. Mitochondria lacking a sustained tubular-to-circular transition were classified as non-contracting.

#### Mitochondrial area quantification

The following quantification was performed in ImageJ/Fiji using the *freehand selection tool*. The total mitochondrial area was quantified in ImageJ/Fiji using the freehand selection tool to trace mitochondria with consistent position and visibility before and after 2-photon laser injury within the imaging ROI (∼28.80 × 28.80 *μ*m; 336 × 336 pixels). Subjects were blinded to experimental conditions using a custom renaming script (renamer.py) developed for this study (see Data Availability for source code). This quantification was used to analyze data for **Figures 1C-1E, S3**.

#### Mitochondrial fold-change quantification

Mitochondrial fold change was quantified by counting the number of mitochondria before and after dendrite injury within the injured branch in Fiji.

#### Mitochondrial ovality index

Quantification was performed in ImageJ/Fiji using the *line tool*. Mitochondrial ovality is an index representing a shape change of individual mitochondria. Mitochondrial minimum and maximum lengths were measured using the line tool in ImageJ/Fiji before and after injury. Ovality before injury was assessed in the intact proximal dendrite by measuring mitochondria prior to the injury frame and at their last visible frame post-injury following primary dendrotomy. Subjects were blinded to experimental conditions using a custom renaming script (renamer.py) developed for this study (see Data Availability for source code).

#### Branching rate and elongation rate quantification

Control neurons and injured neurons were imaged on the day of injury and 24 hrs after injury to generate branching and elongation rates. Neurons were categorized for changes in branching rates based on contracting vs non contracting mitochondria, where contracting groups had at least one contracting mitochondria. Growth rates were calculated by tracing a primary branch and its tertiary branches in an uninjured dendrite of an injured neuron and in control neurons using the Simple Neurite Tracer (SNT) plugin in Fiji. The branching rate calculation is determined by the following equation:

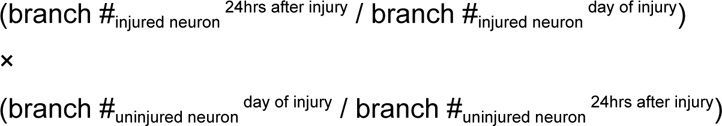

To determine the elongation rate, total dendrite length was used instead of branch #.

#### Mitochondrial shape change analysis

Quantification was performed in ImageJ/Fiji using the *freehand selection tool*. Individual mitochondria were manually traced (ROI) for 30 consecutive frames after first appearing and for the last 30 frames before the end of image acquisition. Changes in mitochondrial area were calculated as the area of a single mitochondria at any frame, divided by the starting area when mitochondria first become visible.

#### Mitochondrial GCaMP analysis

Using the ROI created for tracing the baseline mitoGCaMP intensity (f₀) was defined as the average maximum fluorescence over the first T-10 frames where T = time before mitochondria become visible. Changes in GCaMP fluorescence (Δf/f₀) were calculated as the maximum intensity at any frame divided by f₀. 10 frames before injury, 30 frames immediately after injury, and the last 30 frames were plotted.

### Statistics and reproducibility

Quantification was performed blind for **Figure 1A-1E** and **Figure 4A-4D** using the scrambler code. All statistical analyses were performed in GraphPad Prism 10. The statistical tests used were determined by conducting a normality test on the data distribution. Parametric tests were used if the data followed a normal distribution and non-parametric tests were used if the data did not follow a normal distribution. For parametric tests, if the variance between the compared data were not equal, a Welch’s correction was applied. For non-parametric tests, if the variance between data were not equal, a Mann-Whitney U-test was applied. We conducted ordinary one-way ANOVA for data sets that compared nonparametric versus parametric groups, with *Bonferroni’s* post-hoc correction.

For wildtype vs experimental genotypes, experimental and control genotypes were alternated. Data were only compiled for the final analysis if each independent experiment included at least one experimental and one control animal on the day of data acquisition. Laser injuries were conducted in left or right abdominal segments A1-A6.

## Data Availability

Source data are provided in the supplemental. Code used to blind the users for analysis is deposited in ***GitHub***: https://github.com/pthwu/scrambler. All images used for analysis in this study are deposited in ***Dryad****: All videos and images will be made available upon accepted publication*.

*Raw excel data files used for quantification are attached in supporting information*.

## Acknowledgements

This work was supported by NIH grant R00NS097627 (to KTP) and funding from the Hellman foundation. The authors would also like to acknowledge the University of California Irvine’s Undergraduate Research Opportunities Program and Summer Undergraduate Research Program (UROP and SURP). We would like to thank Dr. Adeela Syed and Dr. Rahul Warrior at the Optical Biology Core at UCI. This study was made possible in part through access to the UCI Optical Biology Core facility of the Developmental Biology Center, a shared resource supported by the Cancer Center Support Grant (CA-62203) and NIH-S10OD032327. We thank members of the Thompson-Peer Lab for critical feedback on the manuscript.

## Funding

**Fig. S1.**
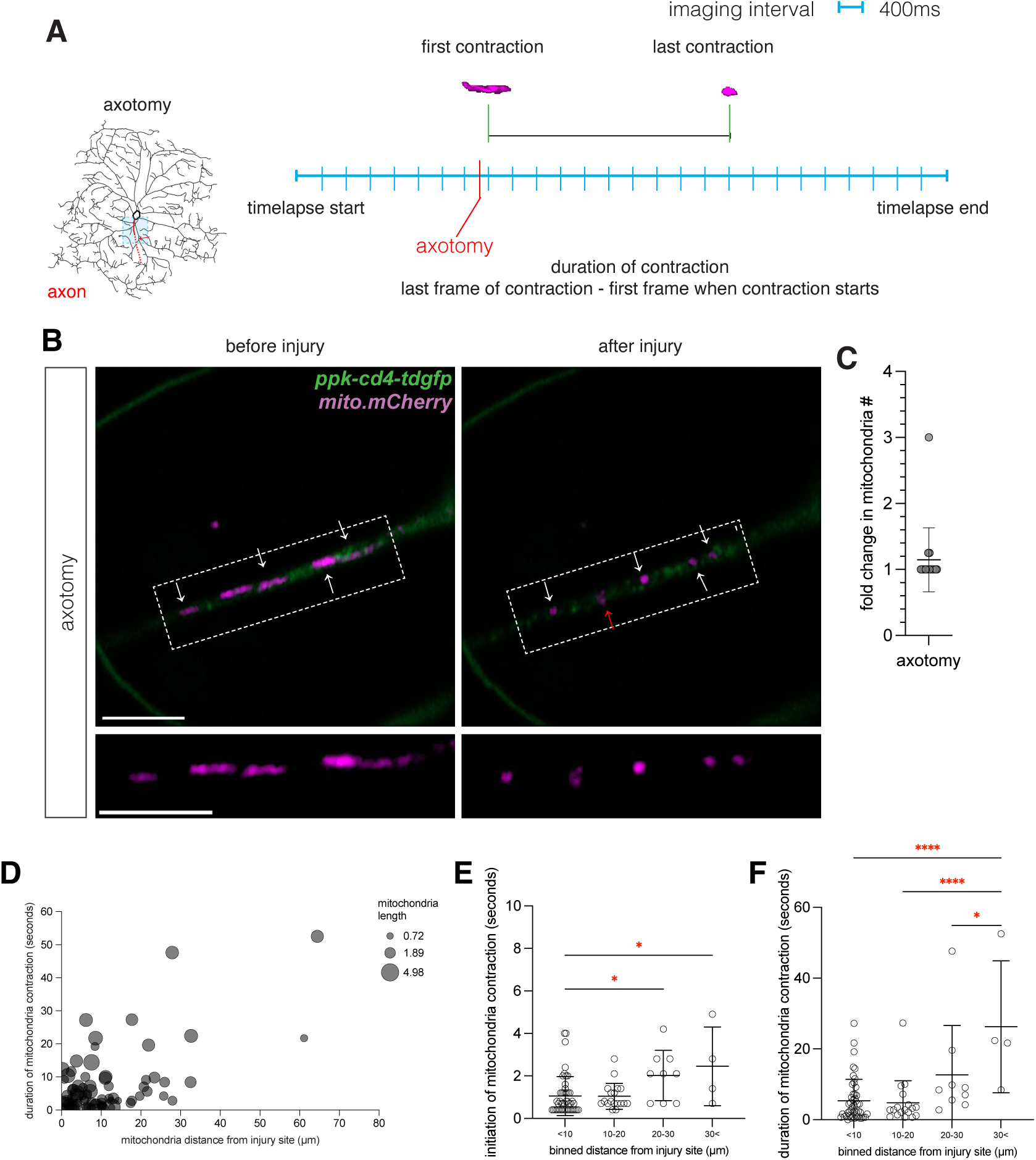
Injury induces mitochondrial contraction in the injured axon. (**A**) Schematic of axotomy and imaging location. (**B**) Representative image of axotomy (red arrow) induced mitochondrial contraction (white arrows) before (left) and after (right) axotomy. Scale bar 5 μm. (**C**) fold change in mitochondria number after axotomy. (**D**) Duration of axonal mitochondrial contraction as a function of distance from axotomy site and mitochondrial length after axotomy in wildtype neurons (N=17, n=78). (**E**) Initiation of axonal mitochondrial contraction after axotomy binned by distance from axotomy site (<10 μm vs. 20-30 μm, p=0.0394*; <10 μm vs. 30< μm, p=0.0371*). (**F**) duration of mitochondrial contraction after axotomy binned by distance from axotomy site. **N** = # of neurons, **n** = # of mitochondria analyzed. wildtype axotomy **N = 17, n = 78** Data presented as mean ± SD. *p<0.05, **p<0.01, ***p <0.001, ****p<0.0001, non-significant p-values are reported. One-way ANOVAs, Bonferroni corrected unless specified.

**Fig. S2.**
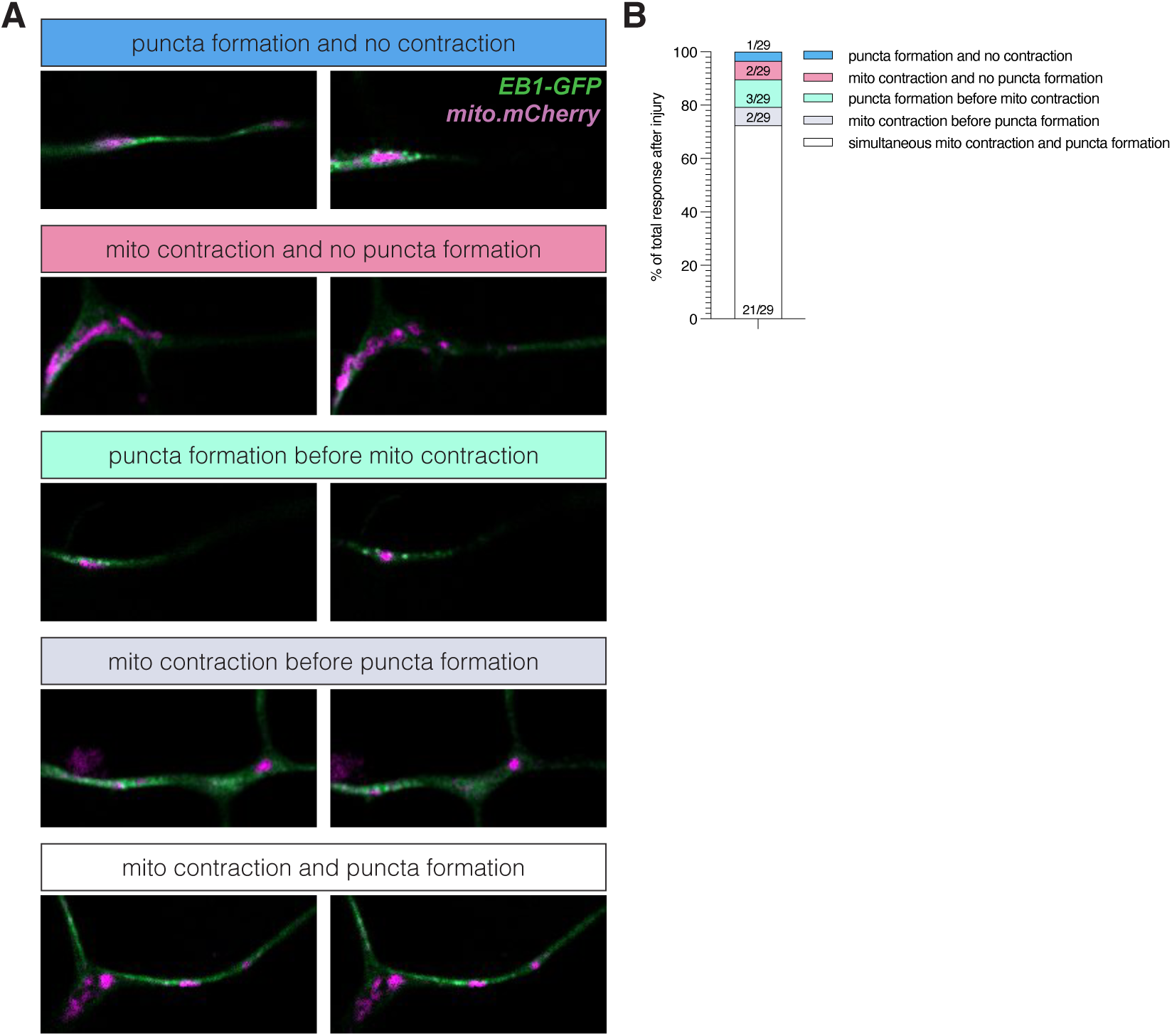
Mitochondrial contraction is not caused by microtubule catastrophe. (**A**) Left columns are before dendrotomy and right columns are after dendrotomy. From top to bottom: microtubules (green, microtubule plus-end binding protein 1, UAS-EB1-GFP) and mitochondria (magenta, UAS-mCherry.mito.OMM) before dendrotomy and the formation of EB1-GFP puncta without mitochondrial contraction 1/29. Mitochondrial contraction in the absence of EB1-GFP puncta formation 2/29. The formation of EB1-GFP puncta before mitochondrial contraction 3/29. Mitochondrial contraction before EB1-GFP puncta formation 2/29. Mitochondrial contraction and EB1-GFP puncta formation occurring simultaneously 21/29. (**B**) Percentage of EB1-GFP mitochondrial contraction outcomes after primary distal dendrotomy. Scale bar 5μm. EB1-GFP distal primary dendrotomy **N = 29**

**Fig. S3.**
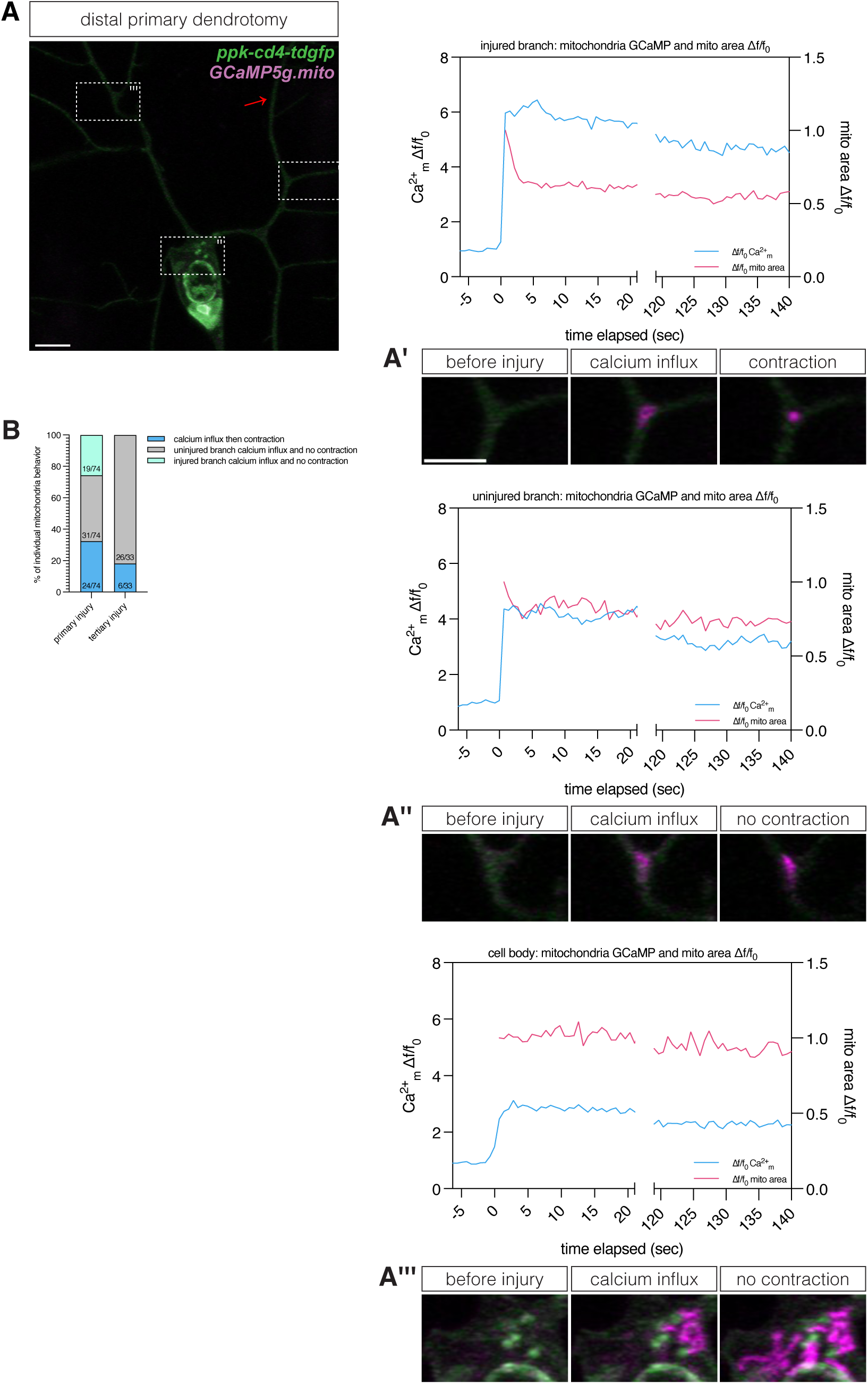
Dendrotomy induces global mitochondrial calcium buffering. (**A**) Mitochondria are not typically visible under baseline conditions with the UAS-GCaMP5g.mito marker but become detectable following dendrotomy (red arrow). Mitochondrial localization (white boxes) in different neuronal compartments post-dendrotomy. N = 7, n = 50. (**A’**) within the injured dendrite, (**A’’**) in the cell body, and (**A’’’**) in an uninjured dendritic branch. (**B**) Quantification of mitochondrial behavior across injured and uninjured dendrites following primary distal dendrotomy (N = 14) and tertiary dendrotomy (N = 7). **N** = # of neurons, **n** = # of mitochondria analyzed. GCaMP5g.mito distal primary dendrotomies **N = 14**; GCaMP5g.mito tertiary dendrotomies **N = 7**

**Fig. S4.**
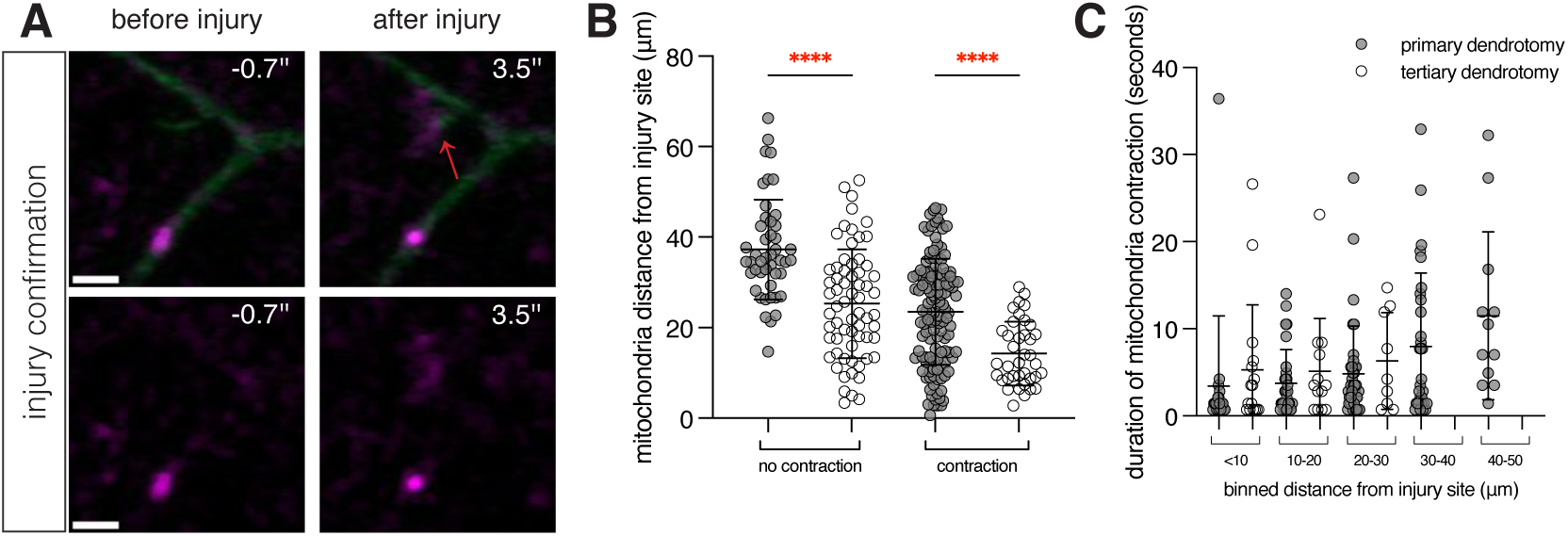
Mitochondrial contraction scales with dendrotomy severity. (**A**) Screenshot of tertiary dendrotomy (red arrow) and mitochondrial contraction. Scale bar 1.5 μm. (**B**) Comparison between average distances away from dendrotomy site that induced no mitochondrial contraction and mitochondrial contraction after distal primary dendrotomy, and tertiary dendrotomy from the neurons analyzed in Figure 2 and Figure 3. One-way ANOVA, Bonferroni corrected. (primary dendrotomy vs. tertiary dendrotomy no contraction, p<0.0001****; primary dendrotomy vs. tertiary dendrotomy contraction, p<0.0001****). (**C**) Binned distance away and duration of mitochondrial contraction for primary dendrotomies vs. tertiary dendrotomies. **Compilation of primary vs tertiary dendrotomies conducted in this study. No additional neurons were injured.** Data presented as mean ± SD.

## Data Transparency

**Fig. 1C** & **1D** was collected from 6 independent experiments across 40 neurons from 21 animals ^ppk-gal4 / ppk-cd4-tdgfp8 ; UAS-mCherry.mito.OMM / +^.

**Fig. 1E** was collected over 3 independent experiments across 22 neurons from 13 animals ^ppk-gal4 / ppk-cd4-tdgfp8 ; UAS-mCherry.mito.OMM / +^.

**Fig. 2D** was collected over 3 independent experiments across 18 neurons from 7 animals ^ppk-gal4 / ppk-cd4-tdgfp8 ; UAS-mCherry.mito.OMM / +^.

**Fig. 2, F–K** was collected over 3 independent experiments across 30 neurons from 14 animals ^ppk-gal4 / ppk-cd4-tdgfp8 ; UAS-mCherry.mito.OMM / +^.

**Fig. 3, C–F** was collected over 10 independent experiments across 36 neurons from 25 animals ^ppk-gal4 / ppk-cd4-tdgfp8 ; UAS-mCherry.mito.OMM / +^.

**Fig. 3, H–L** was collected over 10 independent experiments across 42 neurons from 21 animals ^ppk-gal4 / ppk-cd4-tdgfp8 ; UAS-mCherry.mito.OMM / +^. Out of the 54 animals tested, only 21 were included in the final analysis. Exclusion was based on incomplete ablations between segments and excessive neuron damage (damage to primary branch-See **Video S13** for example), which could compromise data integrity.

**Fig. 4D** was collected over 5 independent experiments across across 13 neurons from 8 animals ^ppk-gal4 / ppk-cd4-tdgfp8 ; UAS-mCherry.mito.OMM / +^ and 21 neurons from 13 animals ^ppk-gal4 / ppk-cd4-tdgfp8 ; UAS-mCherry.mito.OMM / UAS-Hsap|KCNJ2.GFP^. Some of these animals were used to characterize the response in **4E**.

**Fig. 4, G-J** was collected over 8 independent experiments across 46 neurons in 23 animals ^ppk-gal4 / ppk-cd4-tdgfp8 ; UAS-mCherry.mito.OMM / +^ and 46 KCNJ2 neurons in 23 animals ^ppk-gal4 / ppk-cd4-tdgfp8 ; UAS-mCherry.mito.OMM / UAS-Hsap|KCNJ2.GFP^. Out of the 95 animals tested, only 46 were included in the final analysis. Exclusion was based on incomplete ablations between segments, excessive neuron damage (damage to primary branch), and animal death (inability to assess regeneration), poor video quality, which could compromise data integrity.

**Fig. 4K** was part of the 8 independent experiments in **Fig. 4G**, includes larvae that were not able to be assessed for regeneration (**Fig. 4G**) but were able to be assessed for % contraction. **Fig. 5C** was collected over 3 independent experiments across 24 neurons in 11 animals ^ppk-gal4 / ppk-cd4-tdgfp8 ; UAS-mCherry.mito.OMM / +^ and 23 DRP1.K38A neurons in 10 animals ^ppk-gal4 / ppk-cd4-tdgfp8 ; UAS-mCherry.mito.OMM / UAS-DRP1.K38A.HA^.

**Fig. S1** was collected over 6 independent experiments across 25 neurons from 17 animals ^ppk-gal4 / ppk-cd4-tdgfp8 ; UAS-mCherry.mito.OMM / +^.

**Fig. S2** was collected over 3 independent experiments across 29 neurons from 10 animals ^ppk-gal4 / UAS-mCherry.mito.OMM ; UAS-EB1-GFP / UAS-EB1-GFP^.

**Fig. S3A** was collected over 3 independent experiments across 7 neurons from 4 animals ^ppk-gal4 /+ ; ppk-cd4-tdtomato10a / UAS-GCaMP5G.mito^.

**Fig. S3B** was collected over 3 independent experiments across 21 neurons from 5 animals ^ppk-gal4 / + ; ppk-cd4-tdtomato10a / UAS-GCaMP5G.mito^. (Some of this data was used for further analysis for **S3A**).

## Movie Descriptions

**Movie S1.**

ppk-GAL4>UAS-EB1-GFP (homozygote EB1-GFP), 72 hrs AEL, scale bar 5 *μ*m, 10 FPS

**Description**: Axotomy induces microtubule catastrophe, in dendrites and axons, in vivo.

Movie S2.

ppk-CD4-tdGFP[8] ppk-GAL4>UAS-mCherry.mito.OMM, 72 hrs AEL, scale bar 5 *μ*m, 10 FPS

**Description**: Proximal primary dendrotomy induces mitochondrial contraction, in distal severed dendrite, in vivo.

**Movie S3.**

ppk-CD4-tdGFP[8] ppk-GAL4>UAS-mCherry.mito.OMM, 72 hrs AEL, scale bar 5 *μ*m, 10 FPS

**Description**: Distal primary dendrotomy induces local mitochondrial contraction, in proximal intact dendrite, in vivo.

**Movie S4.**

ppk-CD4-tdGFP[8] ppk-GAL4>UAS-mCherry.mito.OMM, 72 hrs AEL, scale bar 5 *μ*m, 10 FPS Description: Axotomy induces mitochondrial contraction, in axons, in vivo.

**Movie S5.**

ppk-GAL4>UAS-EB1-GFP (homozygote) UAS-mCherry.mito.OMM, 72 hrs AEL scale bar 5 *μ*m, 10 FPS

**Description**: Primary dendrotomy induces simultaneous EB1-GFP puncta formation and mitochondrial contraction, in vivo.

**Movie S6.**

ppk-GAL4>UAS-EB1-GFP (homozygote) UAS-mCherry.mito.OMM, 72 hrs AEL scale bar 5 *μ*m, 10 FPS

**Description**: Primary dendrotomy induces mitochondrial contraction before EB1-GFP puncta formation, in vivo.

**Movie S7.**

ppk-GAL4>UAS-EB1-GFP (homozygote) UAS-mCherry.mito.OMM, 72 hrs AEL scale bar 5 *μ*m, 10 FPS

**Description**: Primary dendrotomy induces mitochondrial contraction without EB1-GFP puncta formation, in vivo.

**Movie S8.**

ppk-CD4-tdTomato[10a] ppk-GAL4>UAS-GCaMP5g.mito, 72 hrs AEL, scale bar 5 *μ*m 10 FPS

**Description**: Distal primary dendrotomy induces local mitochondrial contraction, but Ca^2+^ buffering globally, in vivo.

**Movie S9.**

ppk-CD4-tdGFP[8] ppk-GAL4>UAS-mCherry.mito.OMM, 72 hrs AEL, scale bar 5 *μ*m, 10 FPS

**Description**: Tertiary dendrotomy induces local mitochondrial contraction, in vivo.

**Movie S10.**

ppk-CD4-tdGFP[8] ppk-GAL4>UAS-mCherry.mito.OMM, 72 hrs AEL, scale bar 5 *μ*m, 10 FPS

**Description**: Tertiary dendrotomy fails to induce local mitochondrial contraction, in vivo.

**Movie S11.**

ppk-CD4-tdGFP[8] ppk-GAL4>UAS-mCherry.mito.OMM, 72 hrs AEL, scale bar 5 *μ*m, 10 FPS

**Description**: Tertiary dendrotomy induces local mitochondrial contraction, in vivo (Video used for Figure 3h).

**Movie S12.**

ppk-CD4-tdGFP[8] ppk-GAL4>UAS-mCherry.mito.OMM, 72 hrs AEL, scale bar 5 *μ*m, 10 FPS

**Description**: Tertiary dendrotomy fails to induce local mitochondrial contraction, in vivo (Video used for Figure 3i).

**Movie S13.**

ppk-CD4-tdGFP[8] ppk-GAL4>UAS-mCherry.mito.OMM, 72 hrs AEL, scale bar 5 *μ*m, 10 FPS

**Description**: Tertiary dendrotomy induces primary branch degeneration, in vivo.

**Movie S14.**

ppk-CD4-tdGFP[8] ppk-GAL4>UAS-mCherry.mito.OMM / UAS-HSAP\KCNJ2.GFP, 72 hrs AEL, scale bar 5 *μ*m, 5 FPS

**Description**: Injury induced mitochondrial contraction is suppressed by KCNJ2 after primary dendrotomy, in the proximal intact dendrite, in vivo.

**Movie S15.**

ppk-CD4-tdGFP[8] ppk-GAL4>UAS-mCherry.mito.OMM / UAS-HSAP\KCNJ2.GFP, 72 hrs AEL, scale bar 5 *μ*m, 10 FPS

**Description**: Injury induced mitochondrial contraction is partially suppressed by KCNJ2 after primary dendrotomy, in the proximal intact dendrite, in vivo.

**Movie S16.**

ppk-CD4-tdGFP[8] ppk-GAL4>UAS-mCherry.mito.OMM / UAS-HSAP\KCNJ2.GFP, 72 hrs AEL, scale bar 5 *μ*m, 10 FPS

**Description**: Tertiary dendrotomy induces local mitochondrial contraction, in vivo (Video acquired during KCNJ2 replicates).

**Movie S17.**

ppk-CD4-tdGFP[8] ppk-GAL4>UAS-mCherry.mito.OMM / UAS-HSAP\KCNJ2.GFP, 72 hrs AEL, scale bar 5 *μ*m, 10 FPS

**Description**: Tertiary dendrotomy induces local mitochondrial contraction in KCNJ2 expressing neuron, in vivo.

**Movie S18.**

ppk-CD4-tdGFP[8] ppk-GAL4>UAS-mCherry.mito.OMM / UAS-DRP.1K38A.HA, 72hrs AEL, scale bar 5 *μ*m, 10 FPS

**Description**: Loss of *Drp1* inhibits complete mitochondrial contraction after distal primary dendrotomy.

## Notes

Ethical declarations The authors declare no conflicts of interest.

### Competing Interest Statement

The authors have declared no competing interest.

